# Low abundance members of the gut microbiome are potent drivers of immune cell education

**DOI:** 10.1101/2021.11.08.467805

**Authors:** Geongoo Han, Hien Luong, Shipra Vaishnava

## Abstract

One of the main goals of microbiome research is to identify bacterial members that significantly affect host phenotypes and understand their contributions to disease pathogenesis. Studies identifying bacterial members that dictate host phenotype have focused mainly on the dominant members, and the role of low abundance microbes in determining host phenotypes and pathogenesis of diseases remains unexplored. In this study, we compared the gut bacterial community of mice with wide-ranging microbial exposure to determine if low abundance bacteria vary based on microbial exposure or remain consistent. We noted that similar to the high abundance bacterial community, a core community of low abundance bacteria made up a significant portion of the gut microbiome irrespective of microbial exposure. To determine the effect of low abundance bacteria on community structure and host gene expression, we devised a microbiome dilution strategy to “delete” out low abundance bacteria and engrafted the diluted microbiomes into germ-free mice. Our approach successfully excluded low abundance bacteria from small and large intestinal bacterial communities and induced global changes in microbial community structure and composition in the large intestine. Gene expression analysis of intestinal tissue revealed that loss of low abundance bacteria resulted in a drastic reduction in expression of multiple genes involved MHC class II antigen presentation pathway and T-cell cytokine production in the small intestine. The effect of low abundance bacteria on MHC class II expression was found specific to the intestinal epithelium at an early timepoint post-colonization and correlated with bacteria belonging to the family *Erysipelotrichaceae.* We conclude that low abundance bacteria have a significantly higher immuno-stimulatory effect compared to dominant bacteria and are thus potent drivers of early immune education in the gut. Therefore, characterizing immune interaction of low abundance bacteria with the host will offer greater insight into the intestinal immune landscape and disease pathogenesis.

## Background

Bacterial communities living in and on eukaryotic hosts strongly affect host phenotypes including pathogen resistance, inflammation, obesity, behavior, and life span (*1*). Research on the gut microbiome has mainly been restricted to comparisons of the most abundant organisms and the identification of a “core” microbiota associated with health or disease. The core microbiome reflects the capacity of dominant species to exploit the intestinal niche, the available carbon source, nutrients, oxygen level, etc. (*2*). Microbial communities also consist of low abundance bacteria that constitute a significant portion of the microbiome. The “keystone species” concept holds that numerically inconspicuous microorganisms can have an effect on the microbial community and the host that is much greater than their relative abundance (*3, 4*). The “keystone pathogen” concept has been described for several pathobionts that exist in host-associated microbiomes in low numbers but contribute to disease pathogenesis in a major way (*4*). Examples of keystone species that are involved in disease pathogenesis include *Porphyromonas gingivalis* that is associated with periodontitis (*5–9*); *Klebsiella pneumonia*, *Proteus mirabilis* (*10*), and *Citrobacter rodentium* (*11*) associated with intestinal inflammatory diseases; and *Fusobacterium nucleatum* (*12, 13*) associated with colon cancer. Additionally, *Bacteroides fragilis*, a pro-oncogenic bacterium and minor constituent of the colon microbiome in terms of relative abundance can alter colonic epithelial cells and promote oncogenesis due to its unique virulence characteristics (*14*). Collectively these studies demonstrate that studying low-abundant or numerically inconspicuous microorganisms within a microbial community and delineating their effect on community structure and/or host phenotype is crucial for understanding pathogenesis of microbiome-associated complex diseases such as inflammatory bowel disease (IBD).

Experimental manipulations such as removing putative keystone members to assess their impact are used routinely by ecologist studying plants and animal communities (*15, 16*). Such manipulation of keystone members of gut microbiome is challenging owing to difficulty in isolating numerically inconspicuous bacteria many of which have unique growth requirements outside the gut environment. Therefore, empirical evidence showing the impact of the low abundance bacteria on community structure and host phenotype is currently lacking. In this study, we set out to understand the role of low abundance bacteria in stabilizing gut microbial community and their impact on host physiology. We deleted low abundance gut bacteria that were defined as taxa that had a relative abundance of <1% by diluting murine cecal contents and engrafting them into germ-free mice. 16S rRNA gene sequencing and transcriptional analysis of host intestinal tissue of mouse engrafted with microbiomes deleted for low abundance bacteria revealed that low abundance bacteria have a significantly higher immuno-stimulatory effect compared to dominant bacteria and are thus potent drivers of early immune education in the gut. This study thus provides experimental evidence for the key role of low abundance bacteria in host physiology and underscoring the need for studying numerically inconspicuous microbes in disease pathogenesis.

## Materials and methods

### Mice

All mice used were wild-type C57BL/6 background. Female specific-pathogen-free (SPF) mice were purchased from Taconic Biosciences and sacrificed at 8-9 weeks of age. Conventional mice were bred in the SPF barrier facility at Brown University and sacrificed at 7 weeks of age. Female pet store mice were purchased from local pet stores. Germ-free (GF) mice were raised and bred in flexible film isolators gnotobiotic facility at Brown University.

### Cecal microbiota transplantation

Cecal contents were collected from five Taconic mice and suspended in 10 ml PBS followed by filtration with a 70 μm cell strainer. Filtrates were serially diluted tenfold with PBS. Undiluted and serially diluted cecal contents (1:100, 1:1000, and 1:10000 dilution) were stored as a glycerol stock at −80 °C until transplant. 3-4 months old GF mice were orally gavaged with 200 μl of glycerol stock per mouse. Mice were sacrificed after 1- or 5-weeks of transplantation. 4-10 months old wild-type or Het (MyD88^−/+^) GF mice were used for repeated experiments and they were sacrificed after 1-week of transplantation.

### DNA and RNA extraction from the intestine

The small intestine, specifically the ileum, and colon were flushed with PBS and intestinal material was obtained by centrifugation at 5,000 rpm for 10 min. Intestinal contents and fecal samples collected from mice were stored at −80 °C until DNA extraction. Genomic DNA was extracted from ~50 mg of samples using a Quick-DNA Fecal/Soil Microbe Microprep Kit (Zymo Research) according to the manufacturer’s protocol and was stored at −20 °C until further process.

Small pieces of the small intestine and the colon tissue were stored in RNAlater (Invitrogen) at −20 °C until RNA extraction. Total RNA of the small intestine and the colon was extracted using RNeasy Plus Mini Kit (Qiagen) according to the manufacturer’s protocol and was stored at −20 °C until further process.

### Measurement of bacterial load

Bacterial load in the small intestine and the colon was measured by quantitative real-time PCR (qPCR). Reactions were prepared using Maxima SYBR Green/ROX qPCR Master Mix (Thermo Scientific). Bacterial DNA contents in the intestine were determined by 16S rRNA gene contents using the 340F/514R primer pair (340F: 5’-ACTCCTACGGGAGGCAGCAGT-3’, 514R: 5’-ATTACCGCGGCTGCTGGC-3’). Bacterial load was expressed as Ct value and normalized to the weight of starting material.

### 16S rRNA gene sequencing and analysis

The V4/V5 region of the bacterial 16S rRNA gene was amplified from the genomic DNA using the Phusion High-Fidelity DNA polymerase (Thermo Scientific) with 518F/926R primer pair. DNA libraries were constructed as described in our previous study (*17*). The amplicons were sequenced on Illumina MiSeq 2 × 300 bp paired-end sequencing (Rhode Island Genomics and Sequencing Center).

The 16S rRNA gene sequences were processed using QIIME 2 v2019.4 pipeline (*18*). Briefly, sequence reads were denoised and amplicon sequence variants (ASVs) table was produced using DADA2 (*19*). Taxonomic assignment was performed using a pre-trained Naïve Bayes classifier on the SILVA 132 database (*20*). Singletons and all features annotated as mitochondria or chloroplast were removed from the table and the abundance of bacterial taxa was expressed as a percentage of total 16S rRNA gene sequences. For alpha and beta diversity, the feature table was rarefied to even depth. Observed ASVs and Shannon were used as an alpha diversity index. Principal coordinates analysis (PCoA) based on unweighted UniFrac distances was used for beta diversity and differences of sample distances between groups were analyzed using permutational multivariate analysis of variance (PERMANOVA) (*21*). To identify differentially abundant taxa between the groups, ANCOM (*22*) and LEfSe with LDA > 2.0 and p < 0.05 (*23*) were used. R packages phyloseq v1.34.0 (*24*) and qiime2R v0.99.4 (https://github.com/jbisanz/qiime2R) packages were used for visualization. To analyze shared and unique taxa between the groups, R package VennDiagram v1.6.20 was used, and the only taxa that were observed in more than the median number of samples in the group were counted.

### Measurement of gene expression level

Expression of MHC class II-related genes was measured by qPCR. cDNA was synthesized with M-MLV Reverse Transcriptase (Invitrogen). qPCR reactions were prepared using Maxima SYBR Green/ROX qPCR Master Mix (Thermo Scientific) with primer sets for *Ciita* (F: 5’-CGCTGACCTCCCGTGTAAAT-3’, R: 5’-CCTGTCTCTTTAAGAATCGCTCC-3’), *Cd74* (F: 5’-AGAACCTGCAACTGGAGAGC-3’, R: 5’-CAGGCCCAAGGAGCATGTTA-3’), *H2-Aa* (F: 5’-AGGTGAAGACGACATTGAGGAG-3’, R: 5’-GTCTGTGACTGACTTACTATTTCTG-3’) genes. Gene expression was normalized to *Gapdh* and relative expression was calculated using the ΔΔCt method.

### RNA-Seq analysis

Extracted RNA samples were submitted to the GENEWIZ for library construction and sequencing. The RNA library was prepared using NEBNext Ultra RNA Library Prep Kit for Illumina (New England Biolabs) and sequenced on Illumina HiSeq 2 × 150 bp paired-end sequencing (GENEWIZ).

Adapter and low-quality sequences were trimmed from the raw sequence read using Trimmomatic version 0.36 (*25*). Trimmed sequences were aligned to the mm10 mouse genome using STAR v2.7.3a (*26*). Principal component analysis (PCA) and differentially expressed genes (DEGs) analysis were performed in R package DESeq2 v1.30.1 (*27*). Gene ontology (GO) term was analyzed to find enriched pathways using the g:Profiler (*28*). Enriched pathways of the GO biological process (BP) were summarized and visualized with the EnrichmentMap app (*29*) in Cytoscape v3.8.2 (*30*).

### Immunofluorescence staining

Small pieces of the ileum were fixed in formalin for 24 h and embedded in paraffin. Paraffin blocks were sectioned to 7 μm thickness, and slides were deparaffinized with xylenes, ethanol (100%, 95%, and 70%), and water. Antigens were retrieved in a citrate buffer at 95 °C for 20 min. Slides were blocked with 1% bovine serum albumin (BSA) followed by overnight incubation with rabbit anti-EpCAM (CD326) (Invitrogen, cat#MA5-35283) and rat anti-I-A/I-E (Biolegend, cat#107601) antibodies at 4 °C to stain epithelial cells and MHC class II molecules, respectively. After incubation, slides were washed and were incubated with goat anti-rabbit (Invitrogen, cat#A-11008) and goat anti-rat (Invitrogen, cat#A-11081) secondary antibodies for 1 h at room temperature. Slides were counterstained with DAPI and visualized using a Zeiss fluorescence microscope.

MHC class II molecules were quantified using the Analyze Particles function (value of parameters of size and circularity were 0.0001-0.01 and 0.50-1.00, respectively) in ImageJ software (*31*). 6-10 images were used per mouse with 3-4 mice per group.

## Statistical analysis

All statistical analysis and plotting were performed on R v4.0.3 (https://www.R-project.org) and Prism v9.0.2 (GraphPad). Unpaired and two-tailed Mann-Whitney U test or Welch’s t-test was used to find significant differences between the two groups.

## Results

### A core community of low abundance bacteria makes up a significant portion of the gut microbiome

The gut microbiome is composed of a considerable number of bacterial taxa but only a small number of taxa such as the genus *Lactobacillus* and *Bacteroides* take a significant portion of the population. The rest of the population is made up of a lot of low abundance bacteria (*32, 33*). In this study, we first wanted to define the members of high and low abundance bacteria. To achieve this goal, we compared the colon microbiome of SPF mice from Taconic Biosciences (Tac), mice bred and reared in Brown university animal care (Brown), and pet-store mice (Pet store) to understand characteristic features of the high and low abundance members of the gut microbiome in mice with varying environmental exposure to microbes. The gut microbiome of Tac, Brown, and pet store mice consisted of few high abundance genera compared to large number of low abundance bacteria (Fig. S1). Genera that showed higher than 1% in the heatmap (Fig. 1A), were designated as high abundance bacteria while low genera that showed less than 1% of relative abundance were marked as low abundance bacteria. The high abundance bacteria in all three groups were few common genera such as *Lactobacillus*, *Prevotella*, and *Bacteroides* (Fig. 1B). Very few high abundance genera (< 10) accounted for up to 90% of the gut microbiome in the three groups of mice. The high abundance bacteria belonged to only two phyla - *Firmicutes* or *Bacteroidetes* (Fig. 1C and 1D). The low abundance bacteria accounted for only about 10% of the gut bacterial communities, those members consisted of significantly higher number of genera. The low abundance genera belonged to not only *Firmicutes* or *Bacteroidetes* but also various phyla such as *Proteobacteria*, *Actinobacteria*, *Tenericutes*, *Deferribacteres* (Fig. 1C–1E). Next, we assessed if the membership to high and low abundance category in the three microbiomes varied according to the environmental exposure to microbes in the three groups of mice. As expected we saw that the most of the high abundance genera in Tac, Brown, and pet store mice were shared amongst the three group. However, to our surprise we saw that the three groups also shared high proportion of low abundance genera (Fig. 1F). Our results suggest that the low abundance members are the part of core microbiome and that their presence in the gut microbial community is dictated not by environmental exposure but by the host. This finding underscores the need to study the role of low abundance bacteria in regulating gut microbiome community structure and host physiology.

**Fig 1.**
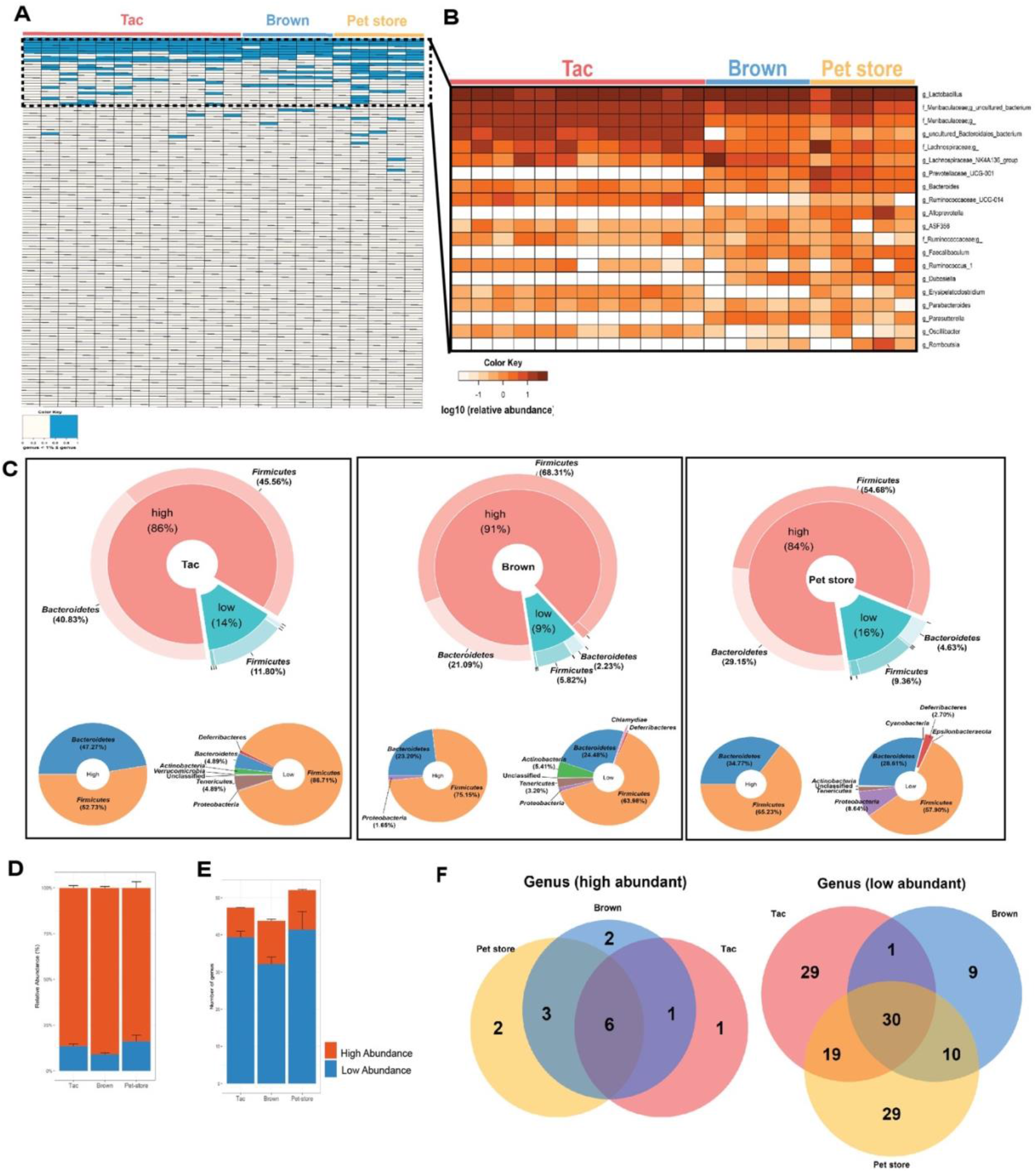
Low abundance bacteria constitute a significant and unique portion of the gut microbiome. (A) Heat map of high and low abundance bacteria based on the fecal 16S rRNA gene of Taconic, Brown, and pet store mice. A genus with a relative abundance of more than 1% is indicated with sky blue. (B) Heat map of 20 most abundance bacteria at the genus level. Donut chart of the phylum composition of high and low abundance bacteria. Relative abundance (D) and the number of the genus (E) of high and low abundance bacteria at the genus level. Error bar represents SEM. (F) Venn diagram of the shared and unique genus of high and low abundance bacteria among the groups.

### Strategy for deleting low abundance bacteria from gut microbiome to study their effect on community structure and host physiology

Bacterial composition in the gut is linked with host physiologies such as metabolic diseases and the immune system (*1, 34, 35*). Low abundance bacteria might contribute to making characteristics of the bacterial community, we tried to explore the role of low abundance bacteria to host in the gut. To exclude a low abundance population from the gut microbiome, we designed an experiment using the dilution method (Fig. 2A). We took cecal contents from Taconic mice and transplanted undiluted and diluted cecal microbiome to the GF mice. At first, we used three different dilution factors (1:100, 1:1000, and 1:10000) to choose an ideal dilution factor to exclude low abundance bacteria from the cecal microbiome with preservation of high abundance bacteria as much as possible. After a week of colonization, we analyzed the colon microbiome of undiluted and three diluted groups by 16S rRNA gene sequencing. In the PCoA plot based on unweighted UniFrac distances, the diluted samples showed a different bacterial composition with undiluted samples and diluted samples were clustered by dilution factor (Fig. S2A). And we looked at their genus composition to compare dilution effects among various dilution factors. In 1:100 dilution, not only most of the high abundance bacteria but also a considerable number of low abundance bacteria were preserved, whereas 1:10000 dilution removed most of the high and low abundance bacteria. Although there was some loss of high abundance bacteria in the 1:1000 dilution, a considerable number of high abundance bacteria survived from dilution and only a few low abundance bacteria were present (Fig. S2B). Based on these results, we chose the 1:1000 dilution factor for further experiments.

**Fig 2.**
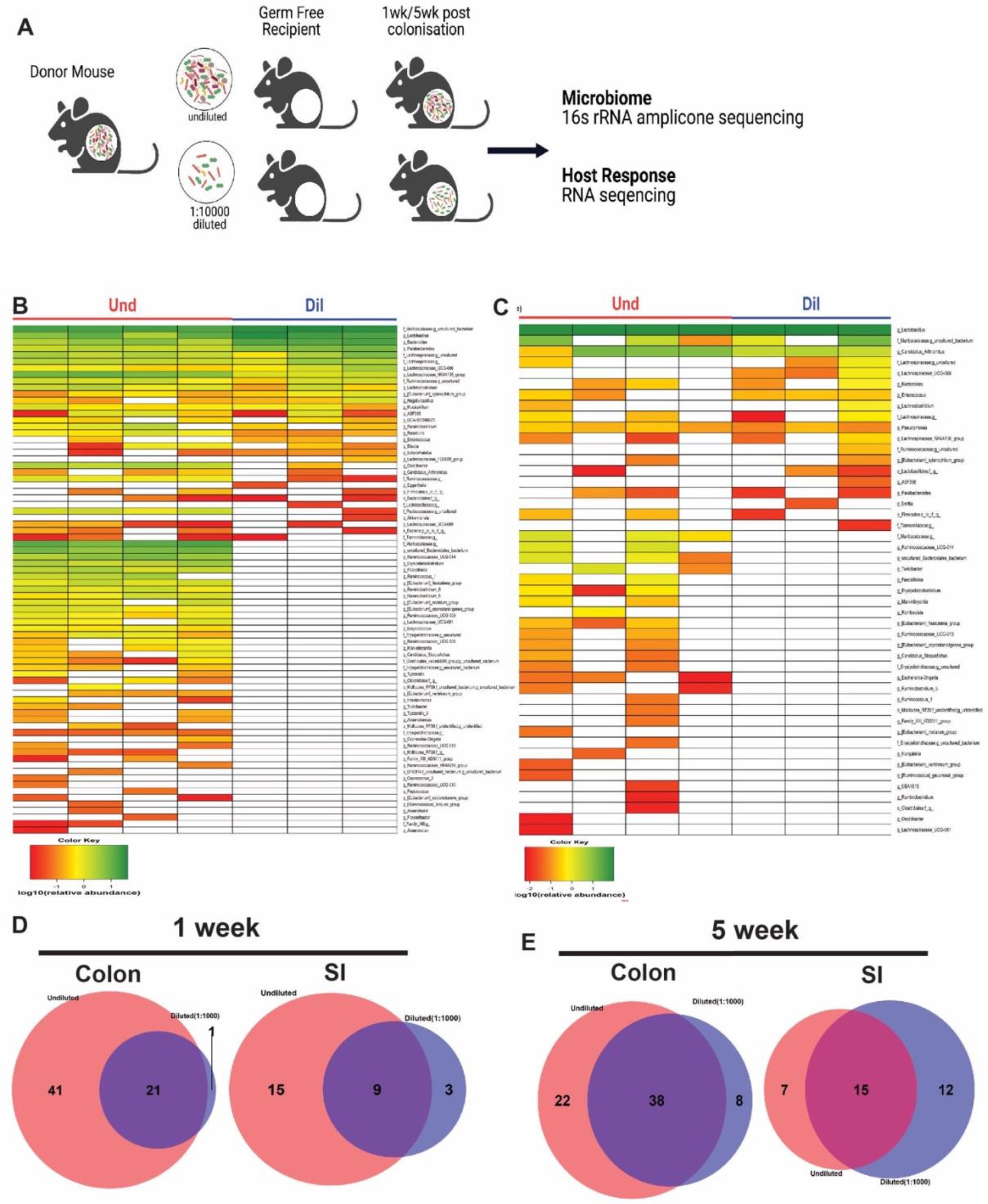
Low abundance bacteria are deleted from the gut microbiome upon colonization of germ-free mice with diluted cecal contents. (A) Strategy for deleting low abundance bacteria from the gut microbiome. Heat map of the colon (B) and small intestine microbiome (C) at the genus level after a week of colonization with the undiluted or diluted microbiome in germ-free mice. In the heat map, a relative abundance of the genus is expressed as log10(relative abundance). The number of shared and unique genera between the Und and Dil group in the colon or small intestine at 1-week post-colonization (D) and 5-week post-colonization (E).

Next, we compared the genus composition between undiluted (Und) and diluted (Dil) groups in the small intestine and colon after one or five weeks of colonization. After a week, the Und and Dil groups shared 21 and 9 genera in the colon and small intestine. A lot of genera were exclusively observed in the Und in both the colon and small intestine (41 and 15 genera, respectively), and most of them were low abundance bacteria (Fig. 2B–2D). In the Dil group, only 1 and 3 genera were exclusively observed in the colon and small intestine, indicating that dilution successfully excluded low abundance bacteria. After five weeks of colonization, however, dilution effects disappeared. In the colon, several genera were still observed in the Und exclusively, but when we compared this result with that of week 1, the number of exclusive genera decreased and many of them were not low abundance genera. The number of unique genera in the diluted group also increased (Fig. S3A and 2E). Effects of dilution also decreased in the small intestine similarly with that of the colon (Fig. S3B and 2E). These results indicate that the mice colonized with Und and Dil over time acquired new genera that were either high or low abundance species.

### Loss of low abundance bacteria from the gut microbiome induces global changes in the bacterial community specifically in the colon

Next, we wanted to assess whether dilution induced global change in the bacterial community structure. 16S rRNA gene amplicon sequencing uses the ASVs, amplicon sequence variants, as the basic unit for analysis (*36*). The ASV is not a taxonomic unit rather it represents each bacterium during marker gene analysis. We compared observed ASVs and Shannon index for alpha diversity (how many taxa are in a sample?) and unweighted UniFrac distances for beta diversity (how many taxa are shared between samples?). In the colon, transplantation of diluted cecal microbiome into the GF mice resulted in lower observed ASVs and Shannon compared to that of the Und group (Fig. 3A) and altered bacterial composition in a week (Fig. 3B). In contrast to the results in the colon, dilution did not reduce diversity and did not alter the composition of the small intestine microbiome. (Fig. 3C and 3D). The interesting thing from these results was the inconsistency between diversity and genus composition, especially in the small intestine.

**Fig 3.**
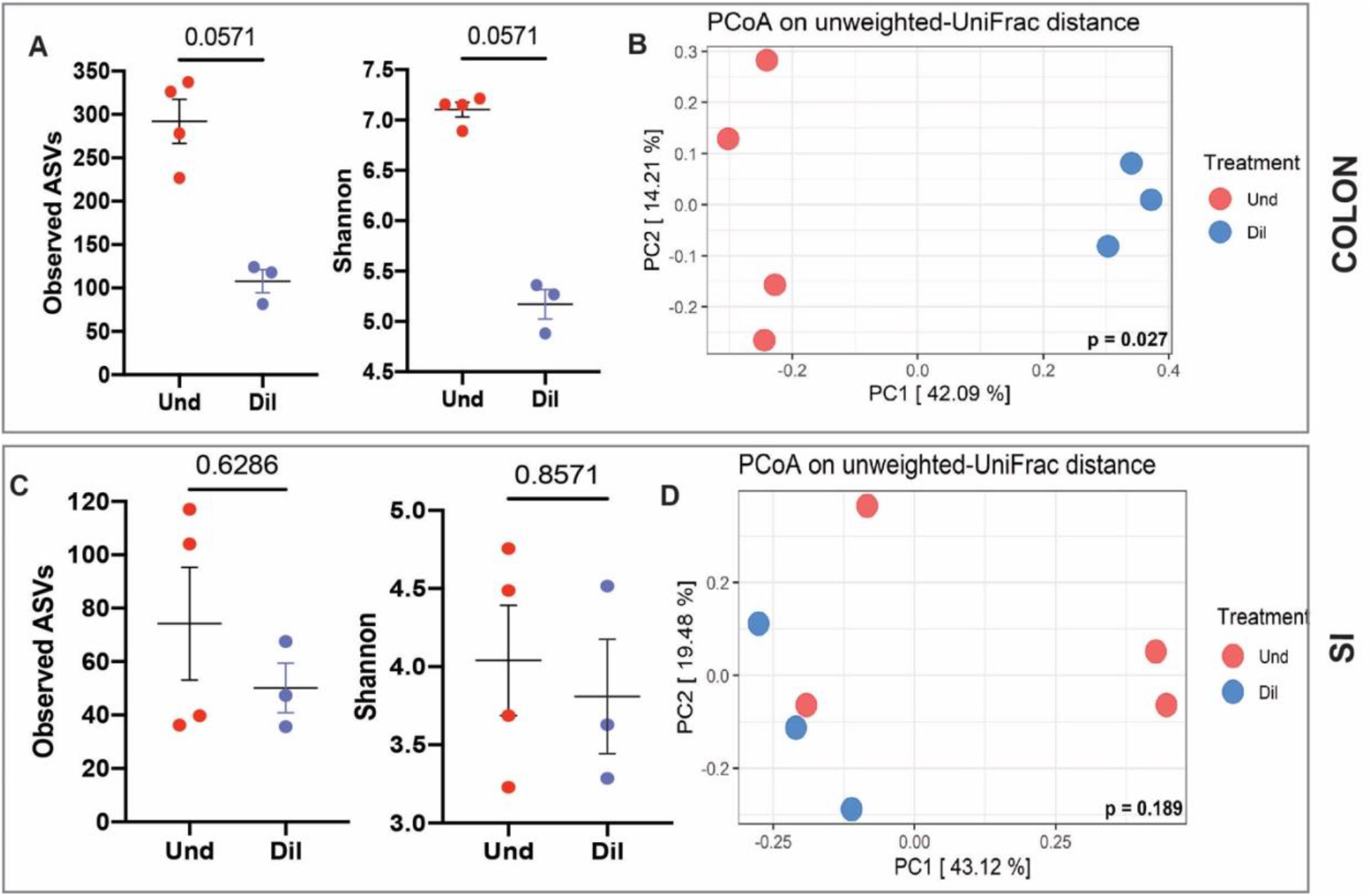
Colonization with diluted cecal contents produced reduced diversity and altered composition at 1 week in the colon bacterial community but not in the small intestine. (A) Alpha diversity (observed ASVs and Shannon) in the colon. (B) PCoA plot based on unweighted-UniFrac distance in the colon. (C) Alpha diversity (observed ASVs and Shannon) in the small intestine. (D) PCoA plot based on unweighted-UniFrac distance in the small intestine. All data were obtained at 1-week post-colonization. Mann-Whitney test and PERMANOVA were used to assess significant differences for alpha and beta diversity, respectively.

Although the Und group had more diverse and exclusive genera than the Dil group, alpha diversity was not significantly different between the groups in the small intestine. Several ASVs can be classified into one or more genera, so the number of ASVs is not directly associated with the number of genera. In addition, some diversity indices such as Shannon consider not only richness but also evenness to get a diversity score. Therefore, we could infer that high abundance genera were composed of more diverse ASVs than low abundance genera and the evenness of ASVs between the two groups was not much different. After five weeks of colonization, similar trends with that of 1-week time point were observed in the colon, whereas the bacterial community was separated into two distinct groups in the small intestine even though diversity was still not different between the Und and Dil (Fig. S4A–S4D). We measured bacterial load in the colon and small intestine and there were no differences between the two groups at 1- and 5-weeks post-colonization (Fig. S5A–S5D) suggesting that engraftment of diluted cecal microbiome into GF mice did not affect bacterial load in the intestine.

### Loss of low abundance bacteria significantly alters expression of MHC class II in the small intestine

To explore the effects of low abundance bacteria on the host, we analyzed gene expression of the small intestine and colon by RNA-Seq. At first, we compared global gene expression between the groups by the PCA. In the PCA plot, the gene expression pattern was clustered into two by treatment in the small intestine, however not in the colon after a week of colonization (Fig. S6A and S6B). After five weeks of colonization, the gene expression pattern was not separated by the group in both the small intestine and colon (Fig. S6C and S6D). Because subjects were clustered into two distinct groups in the small intestine after a week, we analyzed differentially expressed genes (DEGs) to reveal which genes made those differences between the groups. There were 57 up-regulated and 158 down-regulated DEGs in the Und group (Fig. 4A). To infer how those DEGs affect functions in the small intestine, we analyzed the GO term in the context of biological process and several pathways were identified. We drew the enrichment map using the results from GO term analysis for visualization, and similar pathways were clustered together based on the word frequency of the pathway name. Notably, the cluster “Polysaccharide Antigen MHC Class II” and the cluster “Regulation Cytokine Production Process Biosynthetic” were up-regulated in the Und group and these were highly connected with other pathways. Several genes such as *Cd74*, *Cd84*, *Nox1*, *Spn*, and *Ptprc* composed pathways in the cluster “Regulation Cytokine Production Process Biosynthetic”. Pathways in the cluster “Polysaccharide Antigen MHC Class II” were composed of several genes such as *Cd74*, *H2-Aa*, *H2-Ab1*, *H2-DMa*, *H2-DMb1*, *H2-Eb1*, and *March1*, and those genes are related to antigen processing and presentation via MHC class II (Fig. 4B). To focus on the most significant changes by low abundance bacteria at the gene level, we narrowed the list of DEGs down to fifty by FDR, and several genes such as *Cd74*, *H2-Eb1*, *H2-Ab1*, *H2-Aa*, *Ciita*, and *H2-DMa* were up-regulated in the Und group and those are related to MHC class II protein complex or MHC class II transactivator (Fig. 4C).

**Fig 4.**
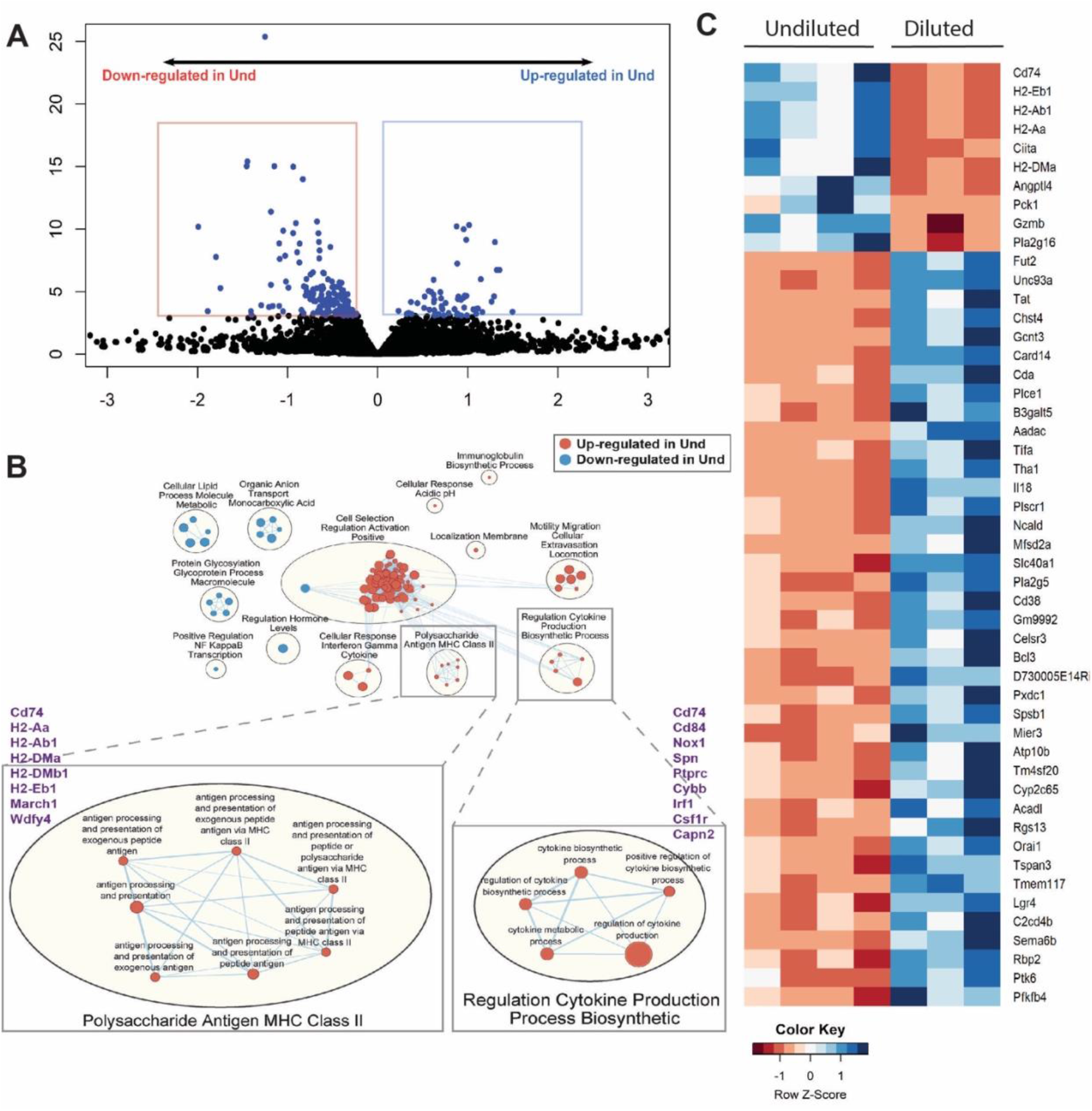
Low abundance bacteria drive the expression of multiple genes involved in antigen presentation and processing pathways in the small intestine. (A) Volcano plot of RNA-Seq results. Each dot represents each gene, and significant DEGs (FDR < 0.05) are expressed as blue dots. Blue and red boxes mean up-regulated and down-regulated DEGs in the Und, respectively. (B) Enriched pathways of the GO biological process. Red and blue circles represent up-regulated and down-regulated pathways in the Und, respectively. (C) Heat map of normalized counts of 50 most significant DEGs. Genes are ordered by FDR. The smaller counts expressed as the redder, the larger counts expressed as the bluer. All data were obtained from the small intestine at 1-week post-colonization.

MHC class II is an essential part of exogenous antigen presentation to the CD4^+^ T cell and is mainly expressed on professional antigen-presenting cells (APCs) but also on intestinal epithelial cells (IECs) (*37, 38*). Several components participate in the MHC class II antigen presentation pathway and the expression of those components is regulated by CIITA in the nucleus. Invariant chain, also known as CD74, stabilizes the MHC class II complex and mediates the assembly and trafficking of that complex (*38*). To assess whether the presence of low abundance bacteria affects the MHC class II antigen presentation, we compared normalized counts of genes that are related to MHC class II, and several important genes for the expression of MHC class II were significantly higher in the undiluted than the diluted group (Fig. 5A). To assess the effects of low abundance bacteria on MHC class II expression at the protein level, we stained the small intestine with the MHC class II marker (I-A/I-E). In the undiluted group, there were more MHC class II molecules than the diluted after a week of colonization, and most of them were only present in the crypts (Fig. 5B and 5C). After five weeks of treatment, however, there was no difference in MHC class II molecules between the two groups, and those molecules were substantially expressed not only in the crypts but also in the villi (Fig. S7).

**Fig 5.**
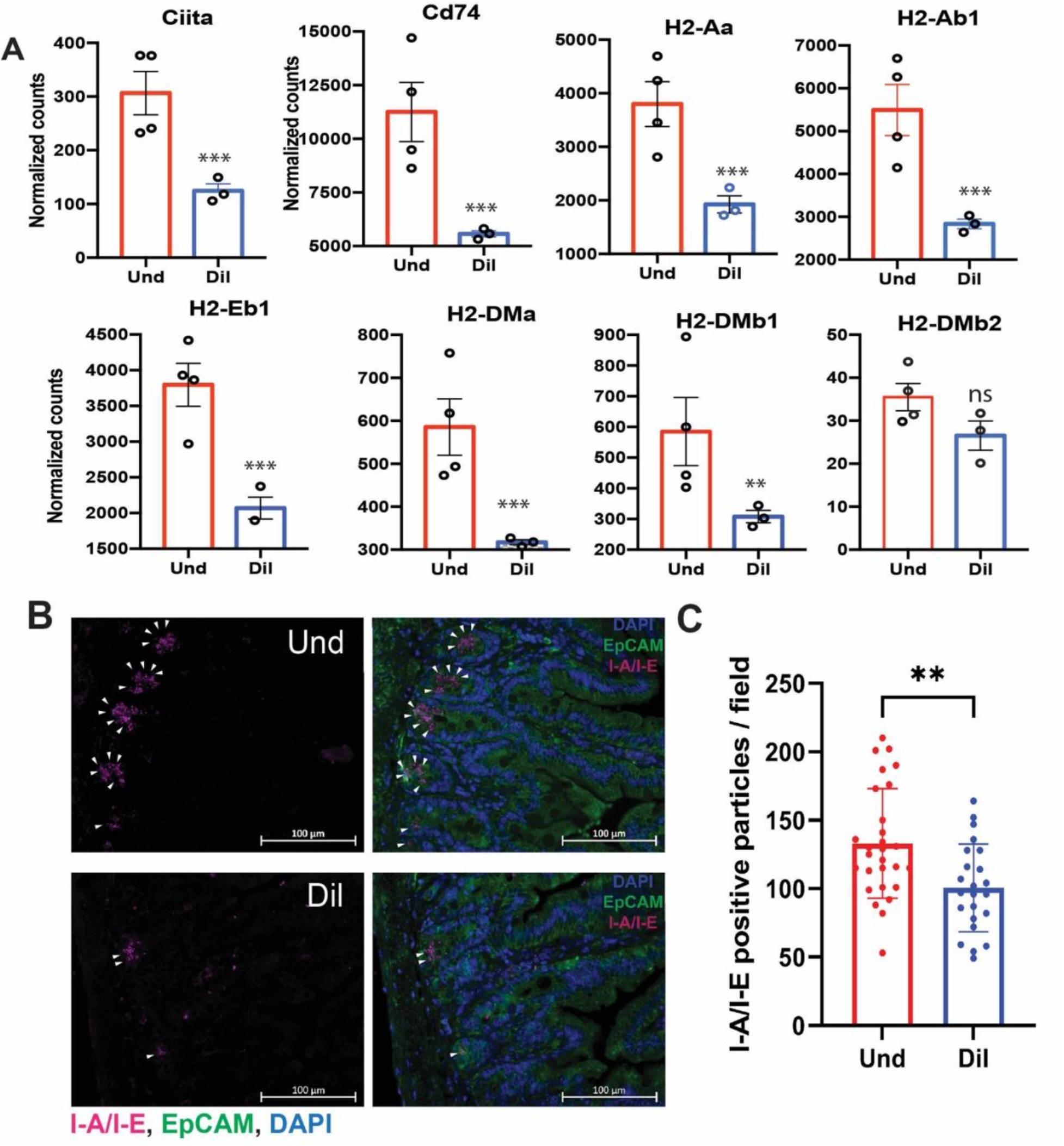
Low abundance bacteria induce MHC class II expression in the small intestine. (A) Normalized counts of MHC class II-related genes in the Und and Dil. DESeq2 was used for statistical analysis. Representative images (B) and quantification (C) of MHC class II expression in the small intestine of the Und and Dil at 1-week post-colonization. Samples were stained with DAPI (nuclei; blue), EpCAM (epithelial cells; green), and I-A/I-E (MHC class II; violet). For quantification of MHC class II molecules, 6-10 images were used per mouse with 3-4 mice per group. Welch’s t-test was used to find significant differences between the two groups.

### Expression of antigen-presenting and processing genes is associated with the presence of low abundance member belonging to family *Erysipelotrichaceae*

Although, a relationship was observed between the expression of MHC class II molecule and low abundance bacteria by RNA-Seq, we wanted to know which member of low abundance bacteria induces expression of antigen presentation and processing genes in the intestinal epithelium. To answer this question, we compared the small intestinal microbiome of mice colonized with undiluted and 1:1000 diluted cecal contents at a 1-week time point and found that there was a significant difference in the relative abundance of bacteria belonging to the family *Erysipelotrichaceae*. In the undiluted group, the relative abundance of the family *Erysipelotrichaceae* was 0.8% but there was no *Erysipelotrichaceae* in the diluted group (Fig. 6A–6C). After 5 weeks of colonization, *Erysipelotrichaceae* was also observed in the diluted group and the relative abundance was not significantly different between the groups (Fig. S8A–S8C).

**Fig 6.**
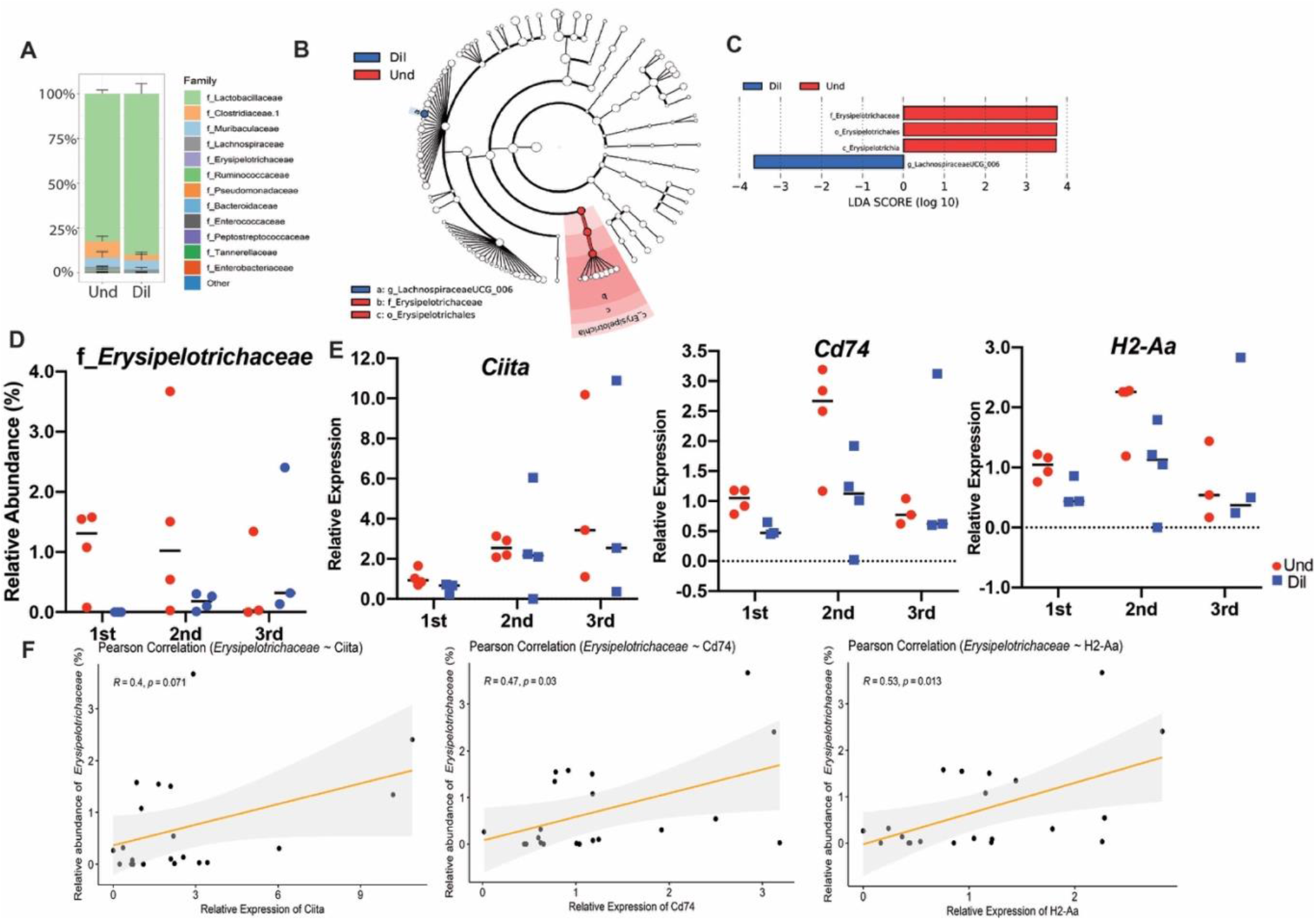
Presence of low abundance bacteria belonging to the family *Erysipelotrichaceae* positively correlates with the enhanced MHC class II antigen processing. (A) Bar plot of relative abundance of the small intestine microbiome at the family level. Error bar represents SEM. Cladogram (B) and bar plot (C), which were obtained from LEfSe analysis, show differentially present taxa between the Und and Dil. Relative abundance of *Erysipelotrichaceae* and relative expression of MHC class II-related genes (*Ciita*, *Cd74*, and *H2-Aa*) (E) in three independent experiments. (F) Scatter plot with regression line to infer the correlation between the *Erysipelotrichaceae* and MHC class II-related genes. Pearson’s correlation coefficients were used to assess the significance of the relationship between bacteria and genes. All data were obtained from the small intestine at 1-week post-colonization.

To be more confident about the role of low abundance bacteria on MHC class II expression in the small intestine, we repeated transplantation of the undiluted and diluted cecal microbiome to GF mice and measured the relative abundance of the *Erysipelotrichaceae* and gene expression level of *Ciita*, *Cd74*, and *H2-Aa*, the representative genes for the expression of MHC class II molecules. Even though we used the same experimental design with the same material as the previous one, the relative abundance of the *Erysipelotrichaceae* and expression of those genes were not higher in the Und compared with that of the Dil group (Fig. 6D and 6E). Because the expression pattern of *Ciita*, *Cd74*, and *H2-Aa* was changed with how abundant the *Erysipelotrichaceae* in each experiment, however, we could strengthen the hypothesis that the *Erysipelotrichaceae* has an important role in the expression of MHC class II in the small intestine. To add more evidence for this hypothesis, we pooled data from three independent experiments and calculated a correlation between that bacteria and each gene, and the *Erysipelotrichaceae* showed strong and significant correlations with *Cd74* (R = 0.47, p = 0.030) and *H2-Aa* (R = 0.53, p = 0.013) although it was not significant with *Ciita* (R = 0.40, p = 0.071) (Fig. 6F).

## Discussion

Skewed pattern in species abundance is a universal feature of ecological communities where there are many rare species but only a few abundant species (*39*). This is also true for host-associated microbial communities such as the gut microbiome where few species dominate the community while numerous species exist in low abundance (*40*). The abundance of a bacteria species in the gut microbiome reflects their ability to exploit the intestinal niche, utilize the available nutrients, and maintain immune homeostasis with the host (*41*). Because abundant bacterial species can be easily identified and isolated their role in the microbial community and their contribution to host physiology are well studied. Abundant species participate in host metabolism via their action on dietary fibers, affect host physiology via their metabolites, regulate immune homeostasis and provide colonization resistance against intestinal pathogen (*42–44*). On the other hand, role of low abundance species in regulating community structure and host physiology remains understudied. Recent work suggests that numerically inconspicuous members of the community can have large effect on community and host despite their low abundance, thus identifying them as keystone members. Examples of keystone microbes that have been linked to disease pathogenesis include *Porphyromonas gingivalis* associated with periodontitis (*6, 45, 46*); *Klebsiella pneumonia*, *Proteus mirabilis* (*10*) and *Citrobacter rodentium* (*11*) associated with intestinal inflammatory diseases; and *Fusobacterium nucleatum* (*12, 13*) associated with colon cancer. Furthermore, studies show that *Bacteroides fragilis* which is a minor constituent of the colon microbiota in terms of relative abundance can alter colonic epithelial cells and mucosal immune function to promote oncogenic mucosal events (*14*). Therefore, studying the minority microorganisms and the nature of their interaction with the hosts could provide novel insights into microbiome-related disease pathogenesis.

There are challenges to studying the low abundance bacteria and their role in host biology. It is hard to differentiate low abundance bacteria that are true colonizers of the gut from the environmental microbes that are just passing by. It is not known if low abundance bacteria form a core community of bacteria or whether their membership in the gut microbiome is accidental, a reflection of hosts’ microbial exposure in a particular environment. Additionally, it is difficult to culture numerically inconspicuous members of the gut community and study how they interact with the host. To differentiate rare bacteria from environmental species we compared abundances of dominant and minority bacteria in the gut microbiome of mice that were exposed to diverse microbes as a virtue of their housing conditions. As expected we saw that irrespective of the microbial exposure (SPF, Barrier or pet store mice), 10 or less genus with relative abundance greater than 1% constituted majority of the microbiome (85%-90%) while about 40 or so genus with relative abundance less than 1% constituted the remaining microbiome. Both high and low abundance bacteria mainly belonged to phylum *Bacteroidetes* and *Firmicutes*, however low abundance bacterial community also included members of phylum *Proteobacteria*, *Tenericutes*, and *Actinobacteria*. We observed that just like dominant bacteria, significant number of low abundance bacteria were shared amongst mice irrespective of their microbial exposure. Presence of core community of low abundance bacteria indicates that they must perform important role in maintaining bacterial community and/or contribute significantly to host physiology thus securing their universal presence in mouse gut microbiome.

To study the role of low abundance bacteria (relative abundance <1%) in regulating community structure and host physiology, we colonized GF mice with cecal contents that were diluted so as to remove majority of low abundance bacteria while keeping high abundance bacteria intact. One week after the engraftment of undiluted and diluted cecal contents 16S rRNA gene analysis revealed that low abundance bacteria were essentially deleted from the bacterial community in the colon as well as small intestine of the mice receiving diluted cecal contents. Additionally, we saw global changes in bacterial community structure with respect to alpha- and beta-diversity in the colon but not in the small intestine of mice that did not harbor low abundance bacteria. The colon presents favorable conditions such as high transit time, optimal pH, low cell turnover and redox potential for the proliferation of bacteria. As a result, colon harbors 70% of the entire gut bacteria and is the major site of bacterial fermentation. Therefore, disruption of community structure specifically in the colon suggests that low abundance bacteria might play a community-specific role in the colon and their deletion might result in more drastic changes in community structure compared to small intestine.

Although small intestine harbors significantly less bacteria than colon, it is a key site for microbiome-induced immune education (*47*). Small intestine resident bacteria directly or indirectly drive differentiation of large number of immune cells that are embedded in intestinal mucosa or within gut-associated lymphoid organs (*48*). Whether immune education in the small intestine happens only in the context of high abundance bacteria as they are the dominant antigens or if low abundance bacteria also provide immune stimuli is not known. To test whether low abundance bacteria drive immune response in the intestinal tissue we analyzed transcriptional response in the small intestine and colon of mice that were deleted for low abundance bacteria and compared it to those that harbored the intact microbiome. We observed that after one week mice that lacked low abundance bacteria had lower expression of multiple genes in MHC class II antigen processing and presentation and cytokine biosynthetic pathways. Reduction in MHC class II marker (I-A/I-E) in mice lacking low abundance bacteria was specifically observed in the crypt cells in the small intestinal epithelium at one week. IECs are capable of MHC class II expression and MHC class II, HLA-DM and invariant chain have been reproducibly detected in IECs throughout all segments of the small intestine (*49–51*). At homeostasis, MHC class II appears to be constitutively expressed on small intestinal enterocytes, mostly in the upper villus (*50, 52*). We saw that at week 5 MHC class II was detected throughout the villi and no difference was observed between mice that lacked low abundance bacteria or had them (Fig. 7). Although it has been known for a while that enterocytes can present antigens, its role in disease pathogenesis or immune homeostasis remains contested. Studies in humans show that MHC class II expression is absent from small intestinal crypts under normal physiologic conditions but is upregulated in specimens obtained from patients with active IBD, celiac disease, and graft vs. host disease (*53–56*). Exposure to inflammatory antigens, such as gliadin in celiac disease, has also been shown to cause the upregulation of cell surface MHC class II and activate effector CD4+ T cells (*54, 57*). Studies in mice however suggest a suppressive role of antigen presentation by IECs, through regulatory T cell activation. Recent studies investigating the role of MHC class II carrying exosomes released by IECs have also reported conflicting findings of either immune enhancing or immunosuppressive activities(*52*). In addition to modulating inflammatory responses, recent findings suggest that MHC class II expression by intestinal stem cells may elicit crosstalk that promotes epithelial renewal. Overall several lines of research in humans and mice suggest an important immunomodulatory role for enterocyte MHC class II. Our correlational studies between most significant microbiome changes and MHC class II expression identified bacteria belonging to *Erysipelotrichaceae* family. Although we cannot rule out involvement of other low abundance microbes in providing the immunostimulatory signals there is evidence for changes in the levels of *Erysipelotrichaceae* in patients with IBD and in animal models of IBD (*58–62*).

**Fig 7.**
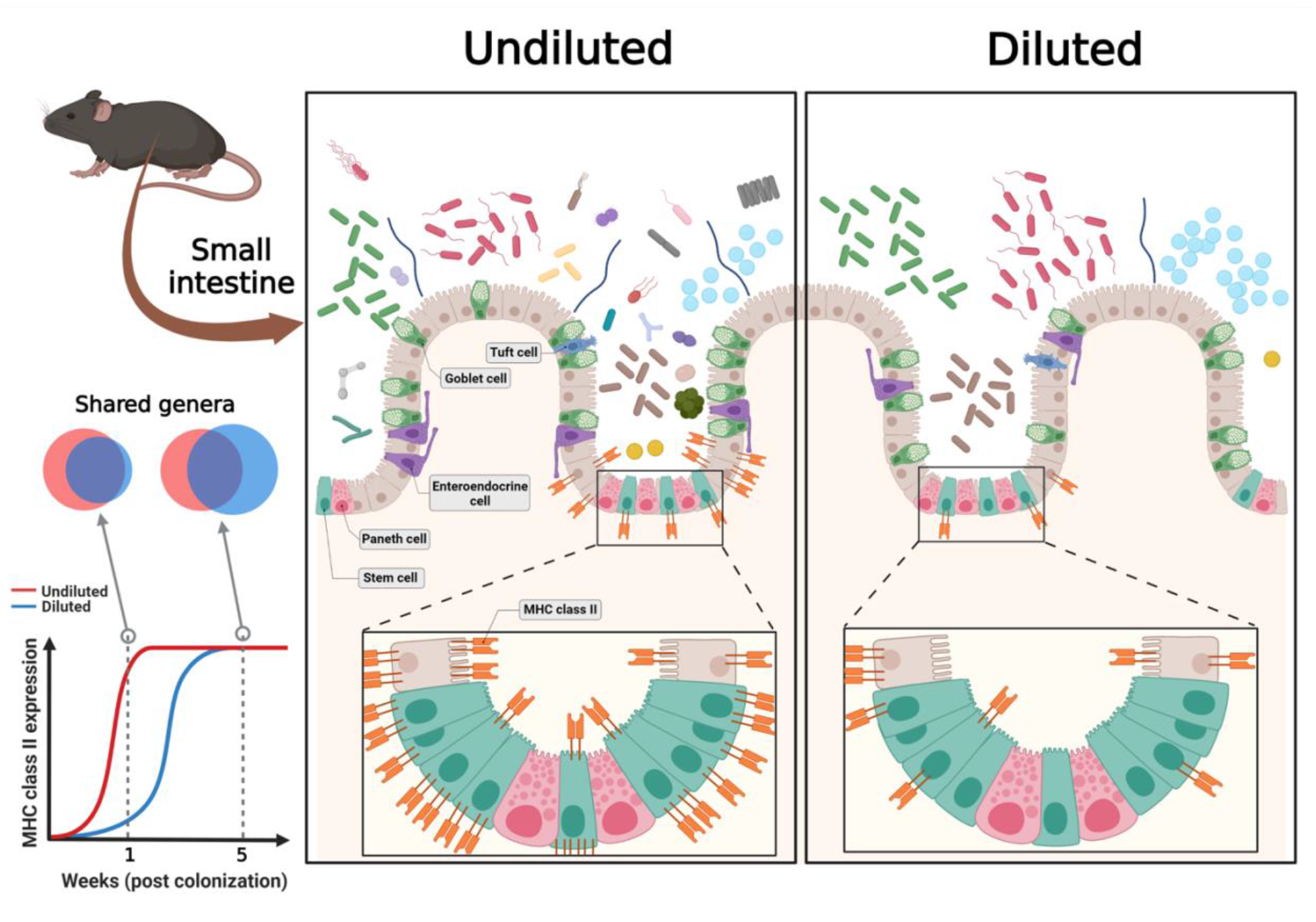
Summary of the interaction between low abundance bacteria and MHC class II expression in the small intestine. Mice gut microbiome consists of a small number of high abundance bacteria with lots of low abundance bacteria. Dilution excludes low abundance bacteria from the cecal microbiome, the diluted microbiome is mostly made up of high abundance bacteria. Colonization of the undiluted and diluted cecal microbiome to the germ-free mice shows that the Und group has almost all genera of the Dil group with exclusive bacterial genera and the different rates of MHC class II expression in the small intestine at earlier time points (week 1; higher expression in the undiluted group than in the diluted group). However, dilution effects on microbial composition disappear and there are no differences of the MHC class II expression at later time points (week 5) (Created with BioRender.com).

## Conclusions

Our work thus provides novel evidence that the low abundance bacteria and not the dominant species are the drivers of inflammatory immune response such as MHC class II expression in the gut and that they should be considered for therapeutic targeting. Although our study mainly focused on low abundance bacteria, other microbes such as viruses and fungi could also drive immune education in the gut. Future studies identifying and assessing the role of novel low abundance microbes in inflammatory disease pathogenesis can be studied using the dilution fecal microbiota transplantation (FMT) approach in animal models and eventually in clinical studies.

## List of abbreviations

APCs: Antigen-presenting cells
ASVs: Amplicon sequence variants
BP: Biological process
BSA: Bovine serum albumin
DEGs: Differentially expressed genes
FMT: Fecal microbiota transplantation
GF: Germ-free
GO: Gene ontology
IBD: Inflammatory bowel disease
IECs: Intestinal epithelial cells
PCA: Principal component analysis
PCoA: Principal coordinates analysis
PERMANOVA: Permutational multivariate analysis of variance
qPCR: Quantitative real-time PCR
SPF: Specific-pathogen-free

## Declarations

### Ethics approval and consent to participate

Experiments were performed according to protocols approved by the Institutional Animal Care and Use Committees of Brown University.

### Consent for publication

Not applicable.

### Availability of data and materials

Raw sequence reads from 16S and RNA sequencing have been deposited in the NCBI Sequence Read Archive (SRA) under accession number PRJNA741381.

### Authors’ contributions

GH and SV designed the study, interpreted the results, and wrote the manuscript. GH performed the experiments. HL managed mice and performed mouse experiments. SV supervised the project. All authors read and approved the final manuscript.

## Acknowledgements

We thank Molecular Pathology Core at Brown University for tissue sample processing. We thank Lalit Beura for providing pet store mice.

## Supplementary Information

**Fig. S1.**
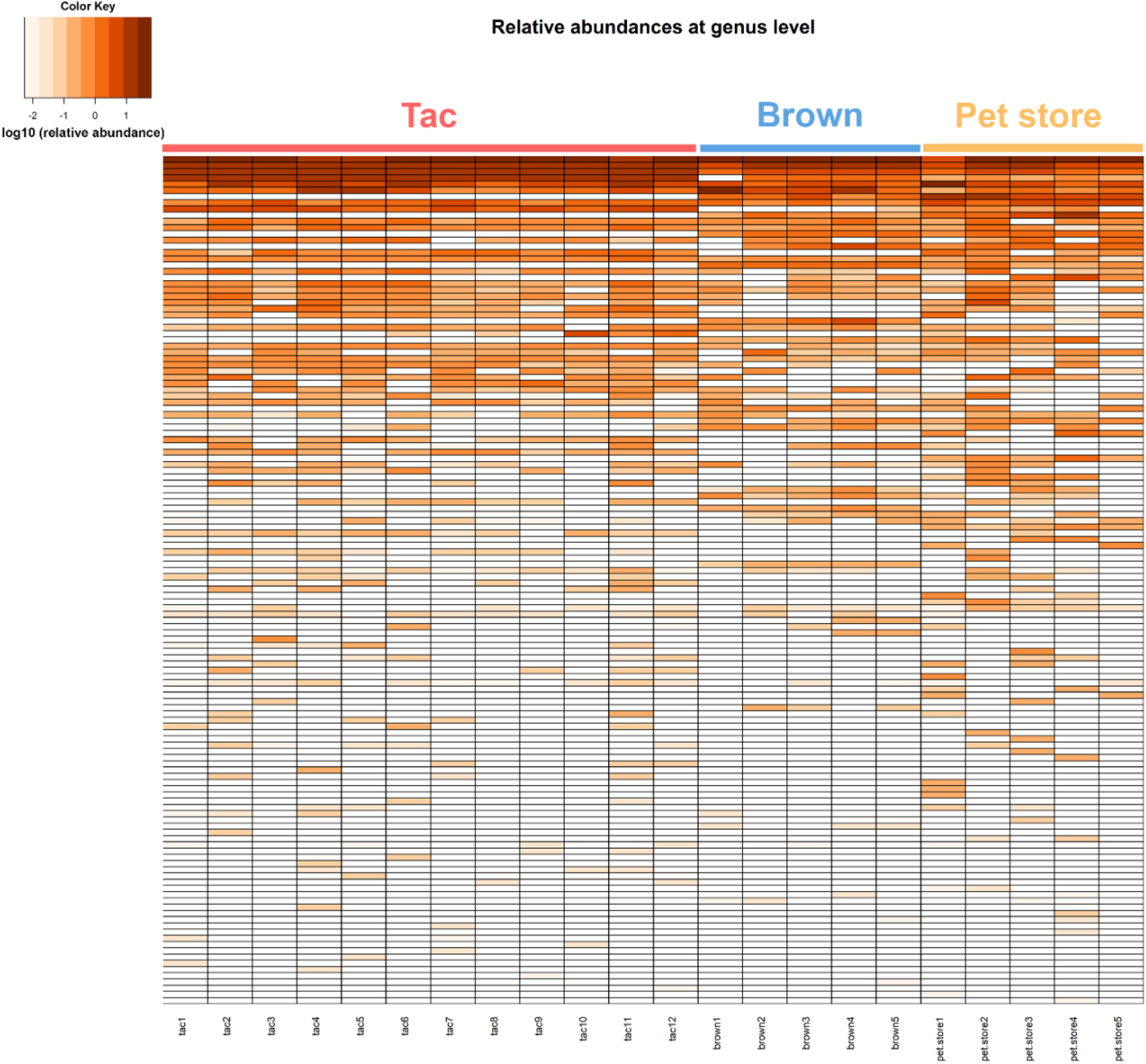
Heat map of relative abundance at the genus level of the gut microbiome of Taconic, Brown, and pet store mice.

**Fig. S2.**
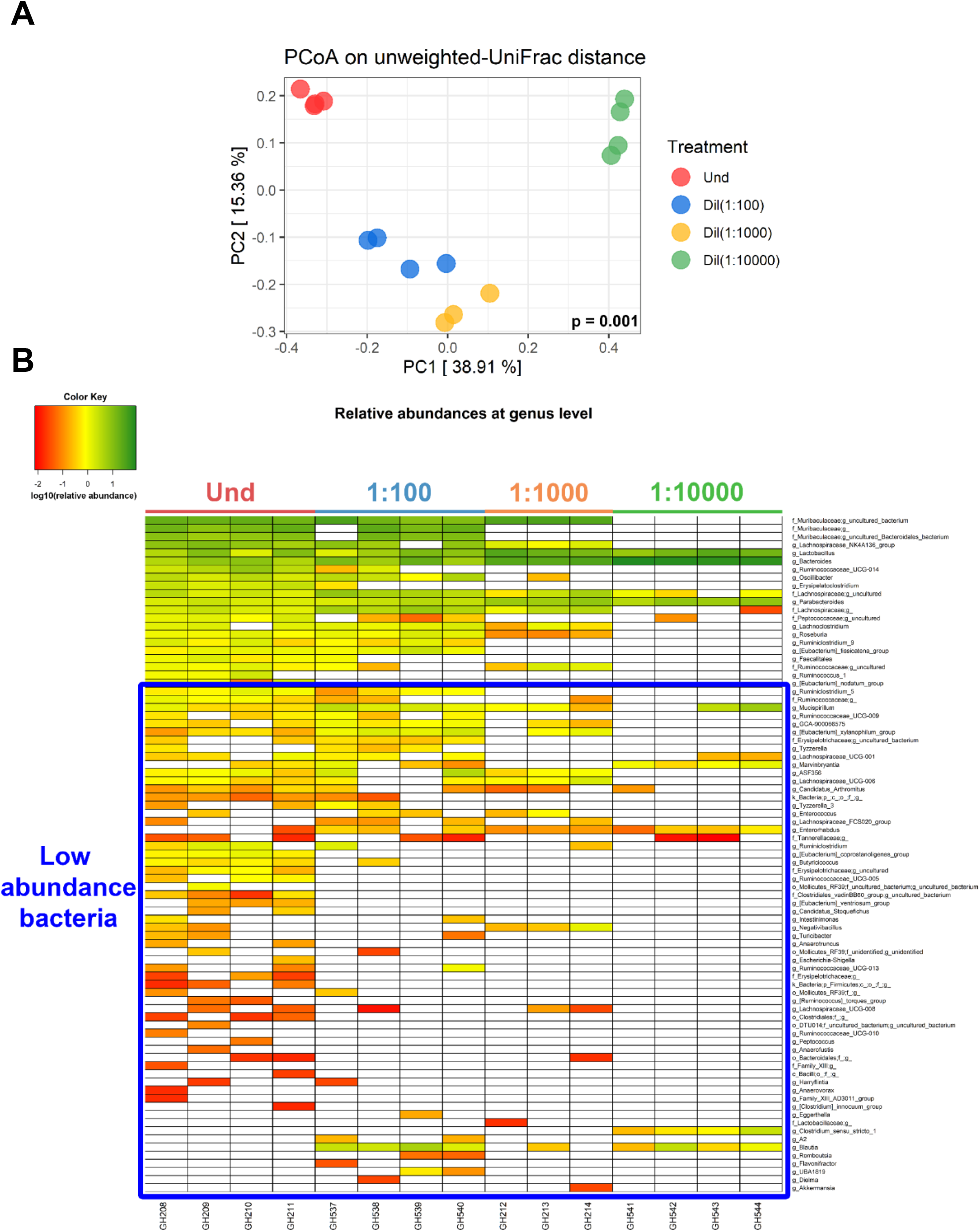
Composition of the colon microbiome in germ-free mice colonized with diluted cecal contents at various levels of dilution. (A) PCoA plot based on unweighted-UniFrac distance. (B) Heat map of the relative abundance at the genus level. Blue box represents low abundance bacteria of the undiluted cecal contents. A relative abundance of the genus is expressed as log10(relative abundance).

**Fig. S3.**
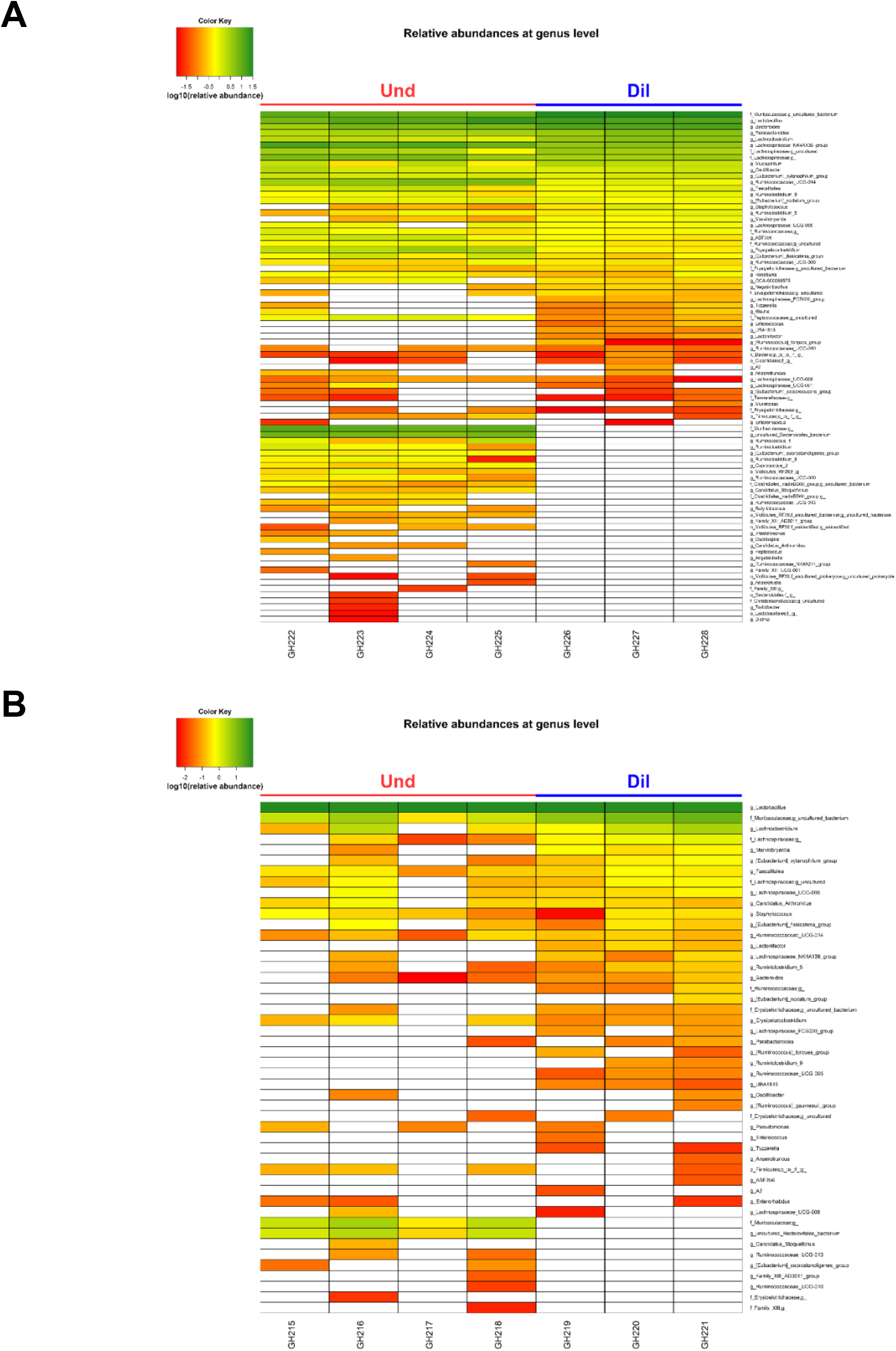
Heat map of the colon and small intestine microbiome at the genus level after five weeks of colonization with the undiluted or diluted microbiome in germ-free mice. (A) Colon. (B) Small intestine. A relative abundance of the genus is expressed as log10(relative abundance).

**Fig. S4.**
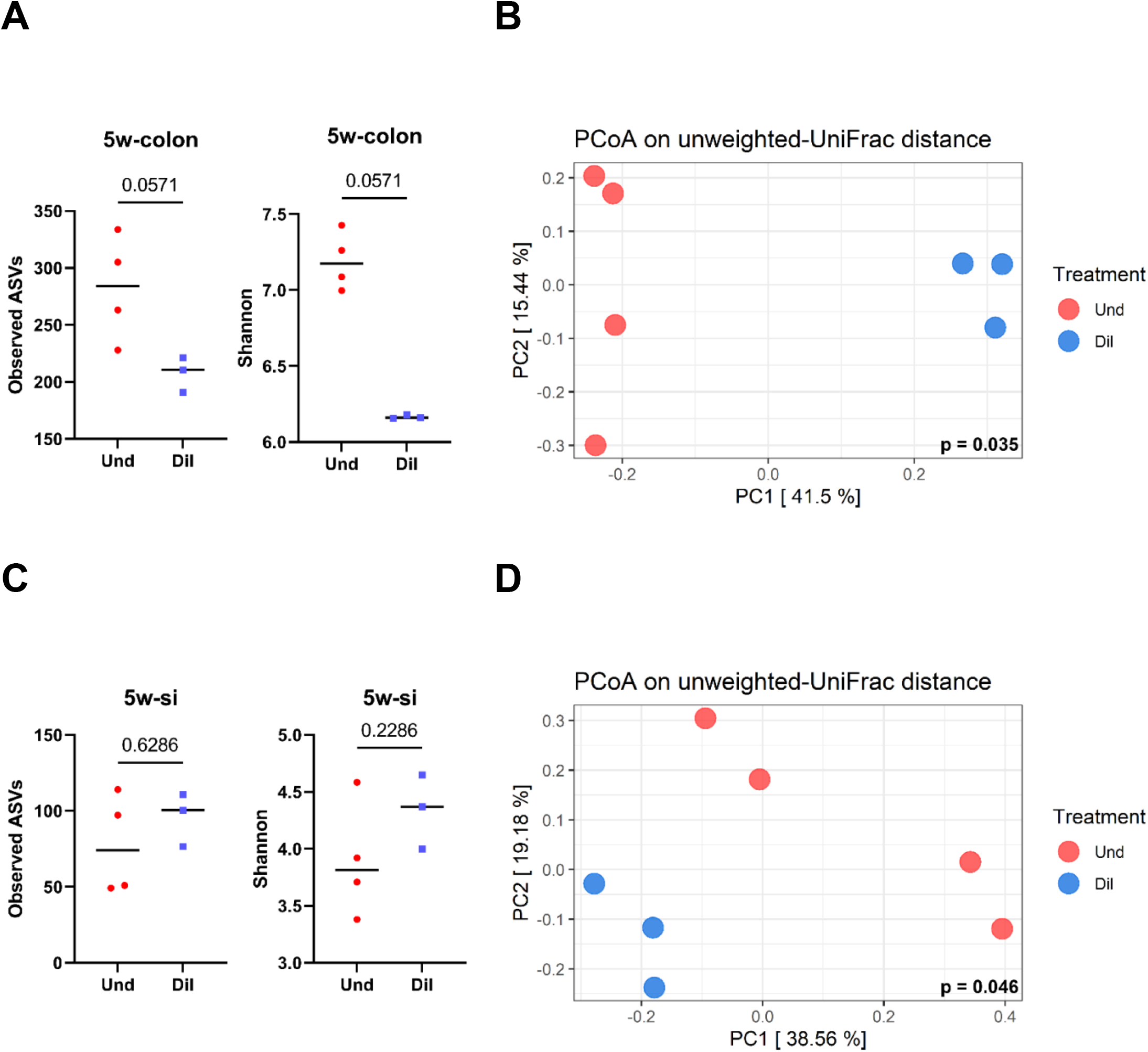
Alpha and beta diversity of the Und and Dil group in the colon and small intestine at 5-weeks post-colonization. (A) Alpha diversity (observed ASVs and Shannon) in the colon. (B) PCoA plot based on unweighted-UniFrac distance in the colon. (C) Alpha diversity (observed ASVs and Shannon) in the small intestine. (D) PCoA plot based on unweighted-UniFrac distance in the small intestine. Mann-Whitney test and PERMANOVA were used to assess significant differences for alpha and beta diversity, respectively.

**Fig. S5.**
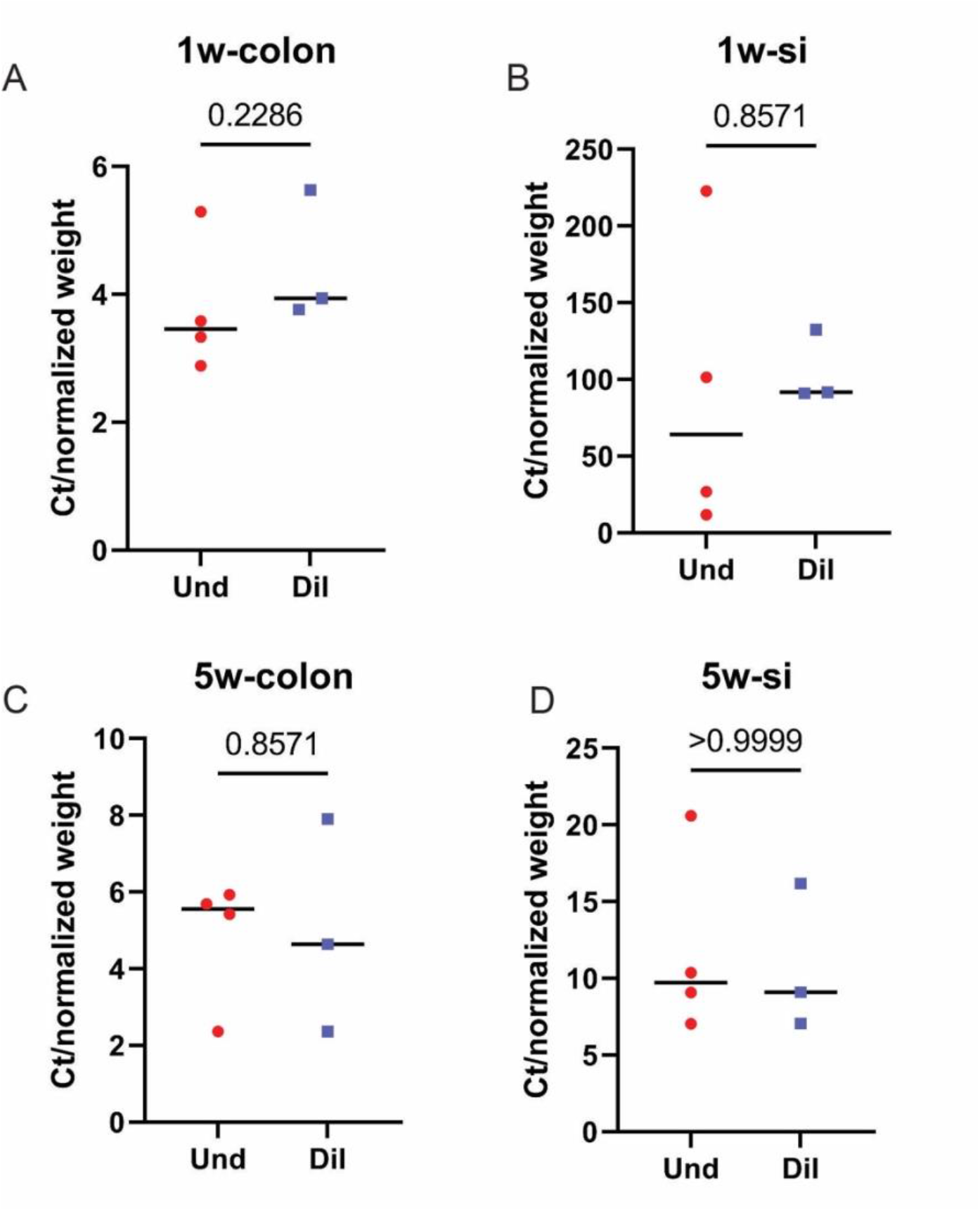
Bacterial load in the colon and small intestine at 1- and 5-weeks post-colonization. Bacterial load in the colon (A) and small intestine (B) at 1-week post-colonization. Bacterial load in the colon (C) and small intestine (D) at 5-weeks post-colonization. Mann-Whitney test was used to assess significant differences between the two groups.

**Fig. S6.**
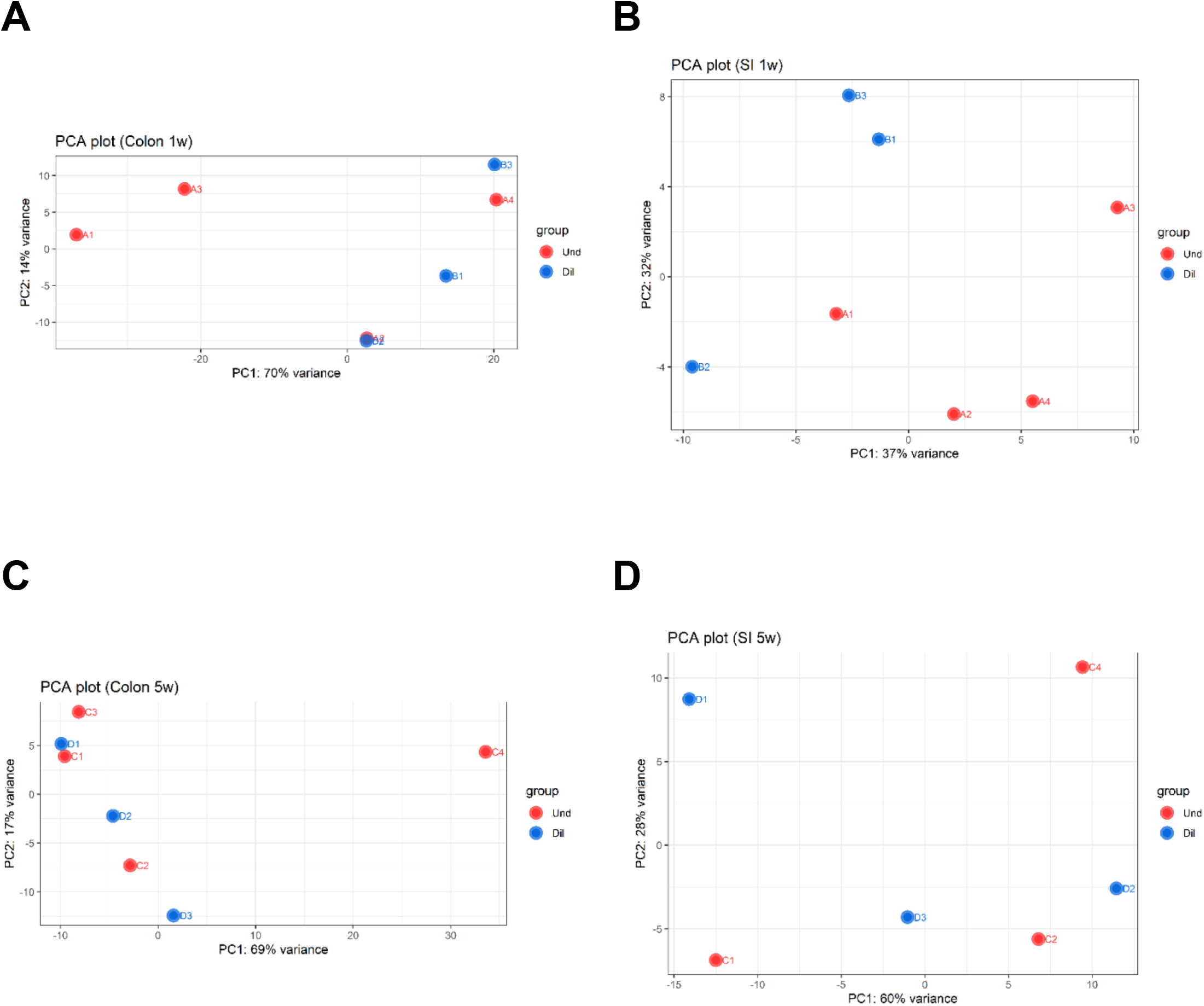
Global gene expression pattern based on RNA-Seq results. PCA plot in the colon (A) and small intestine (B) at 1-week post-colonization. PCA plot in the colon (C) and small intestine (D) at 5-weeks post-colonization.

**Fig. S7.**
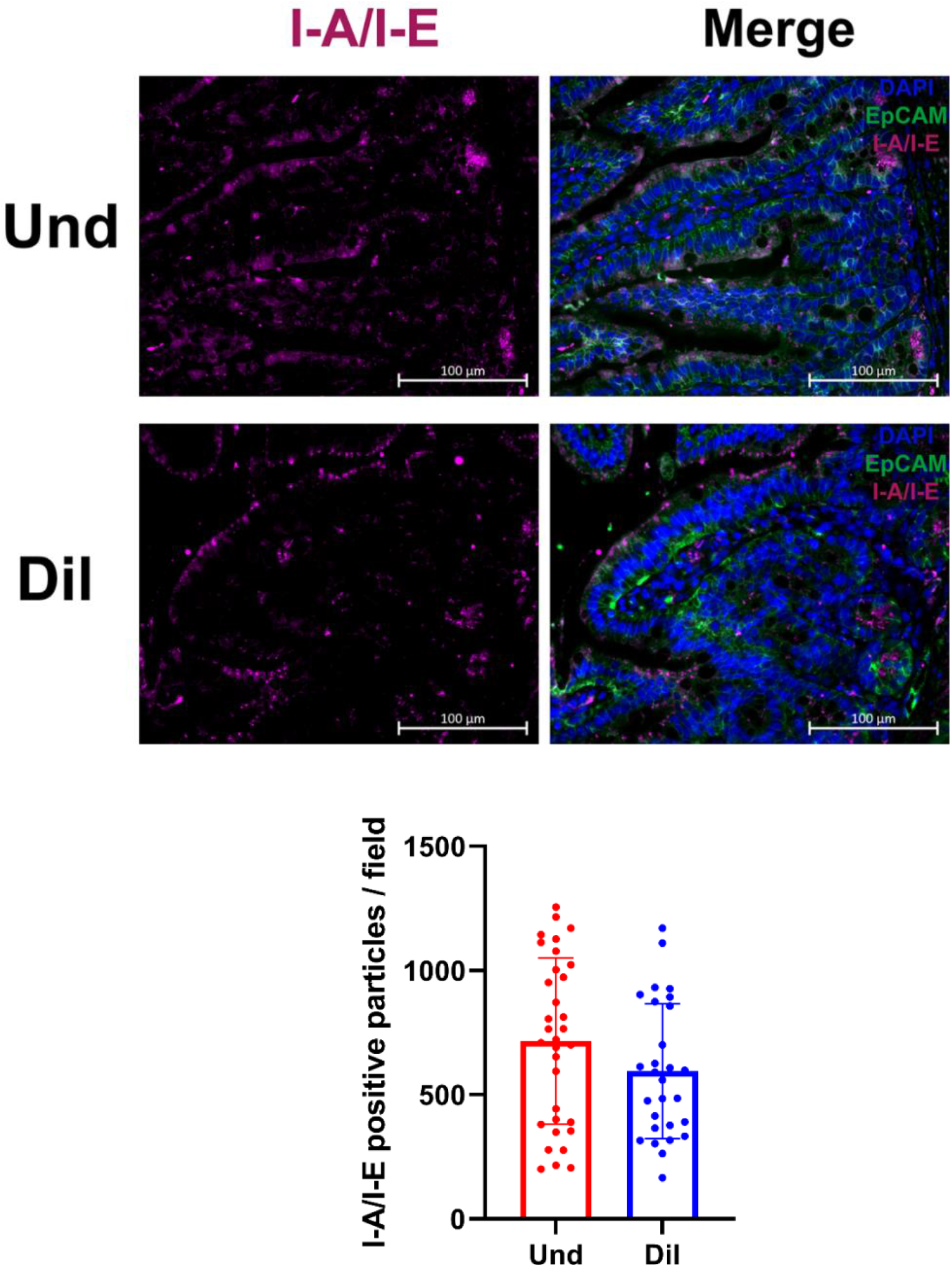
Representative images and quantification of MHC class II expression in the small intestine of the Und and Dil at 5-weeks post-colonization. Samples were stained with DAPI(nuclei; blue), EpCAM (epithelial cells; green), and I-A/I-E (MHC class II; violet). For quantification of MHC class II molecules, 6-10 images were used per mouse with 3-4 mice per group. Welch’s t-test was used to find significant differences between the two groups.

**Fig. S8.**
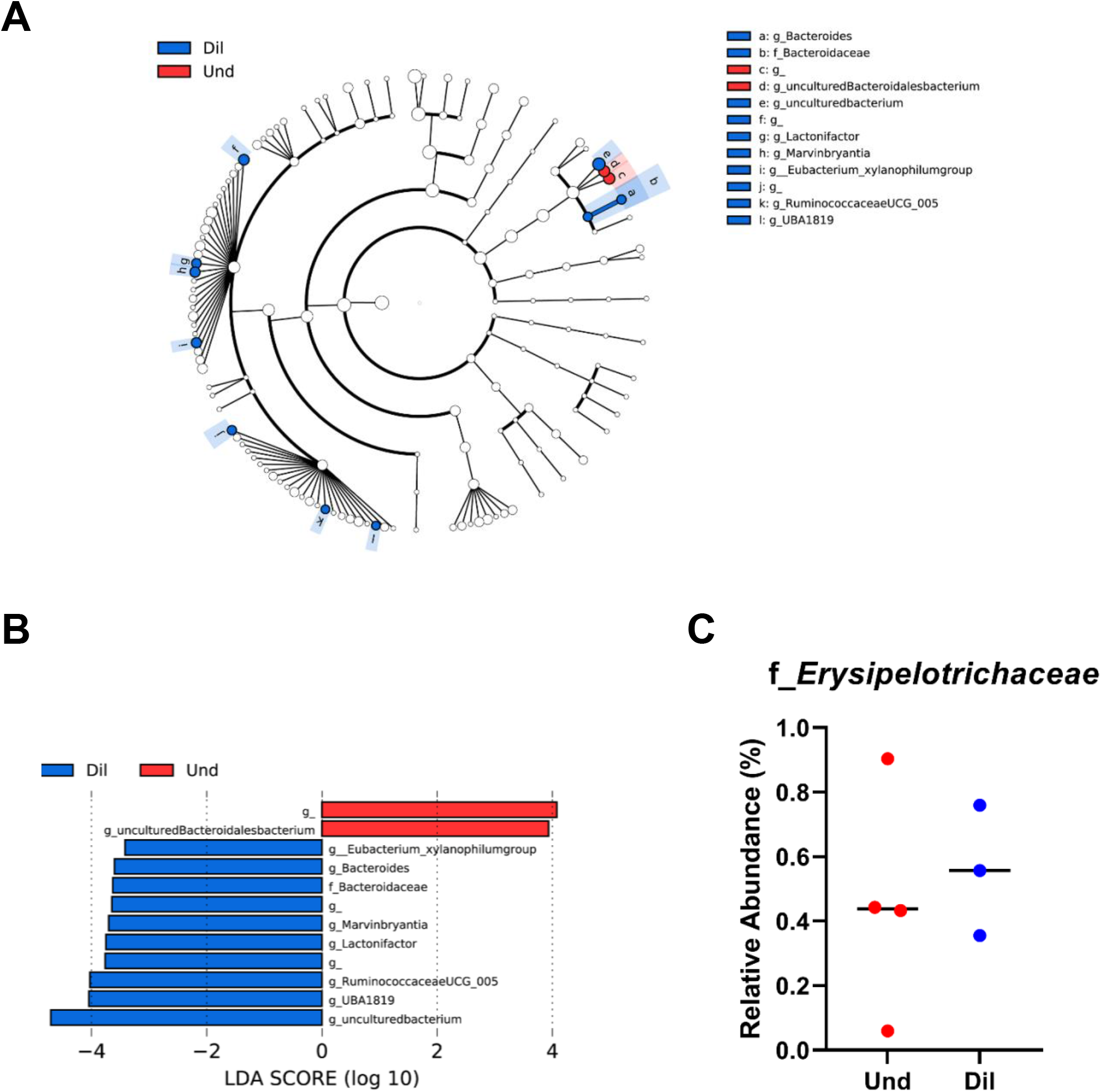
Differentially abundant taxa between the Und and Dil in the small intestine at 5 weeks post-colonization. Cladogram (A) and bar plot (B), which were obtained from LEfSe analysis, show differentially present taxa between the Und and Dil. (C) Relative abundance of *Erysipelotrichaceae*.

